# Large genetic divergence underpins cryptic local adaptation across ecological and evolutionary gradients

**DOI:** 10.1101/2021.12.05.471216

**Authors:** Morgan M. Sparks, Joshua C. Kraft, Kliffi M. Subida Blackstone, Gordon G. McNickle, Mark R. Christie

## Abstract

Environmentally covarying local adaptation is a form of cryptic local adaptation in which the covariance of the genetic and environmental effects on a phenotype obscures the divergence between locally adapted genotypes. Here, we systematically document the magnitude and drivers of the genetic effect (V_G_) for two forms of environmentally covarying local adaptation: counter- and cogradient variation. Using a hierarchical Bayesian meta-analysis, we calculated the overall effect size of V_G_ as 1.05 and 2.13 for populations exhibiting countergradient or cogradient variation, respectively. These results indicate that the genetic contribution to phenotypic variation represents a 1.05 to 2.13 standard deviation change in trait value between the most disparate populations depending on if populations are expressing counter- or cogradient variation. We also found that while there was substantial variance among abiotic and biotic covariates, the covariates with the largest mean effects were temperature (2.41) and gamete size (2.81). Our results demonstrate the pervasiveness and large genetic effects underlying environmentally covarying local adaptation in wild populations and highlights the importance of accounting for these effects in future studies.

## Introduction

How and when populations can persist via adaptation to local environmental conditions remains a central question in ecology and evolutionary biology [1]. Environmental gradients, where abiotic and biotic factors such as temperature, salinity, or predator type vary in a predictable manner, provide a framework to formulate and test questions about the pattern and magnitude of local adaptation. Pairing environmental gradients with common garden or reciprocal transplant studies, especially when there are many populations distributed along these gradients, can lead to insights into how local adaptation covaries with specific environmental factors [2,3].

Shared patterns of phenotypic variation along ecological gradients spurred notable research on the environmental and genetic determinants of body size and morphology and led to the establishment of eco-geographical “rules” such as Bergmann’s, Allen’s, and Hesse’s rules and subsequent derivations therein [4-9]. Foundational work by Levins [2,10] revealed a pattern whereby fruit flies distributed along an elevation gradient expressed a similar body size in their wild home environments, leading to the conclusion that the trait was canalized. However, a subsequent study revealed the higher elevation populations were much smaller than lowland populations in common environments—expressing disparate phenotypes across all common environments, a response believed to be related to desiccation—suggesting these populations were locally adapted to different elevations. In concert, these findings provided the first formal evidence of environmentally covarying local adaptation.

The variance components of phenotypic plasticity are commonly expressed with reaction norm plots, where the trait value of a genotype is plotted as a function of multiple environmental values [11]. In a reaction norm, the slope of a genotype’s response is representative of phenotypic plasticity (the effect of the environment) and the difference between genotype-specific reaction norms at each environmental variable is representative of the genotypic difference (the genetic effect). When different genotypes express the same reaction norm (*i.e*., overlapping lines, Fig. 1a, plasticity alone) this is representative of a purely environmentally driven response where distinct genotypes would express the same trait values (or phenotype) along an environmental gradient. Alternatively, populations may be locally adapted, such that the reaction norms—namely, fitness or fitness related traits—cross when comparing populations in their respective home and away environments, which is indicative of a genotype by environment interaction (Fig. 1b, Genetic x Environment). Finally, populations may exhibit a form of cryptic local adaptation, where the genetic and environmental effects covary with one another, either negatively or positively [12]. In this scenario, the reaction norms for populations may have the same or similar slopes thus maintaining consistent rank order in their trait values (Fig.1, co- and countergradient variation). For example, a high latitude population may have consistently larger body sizes across environments than a low latitude population, despite both expressing plasticity in body size. When the genetic portion of phenotypic variance opposes the environmental portion, it is called countergradient variation (negative covariance; Fig. 1c), and when those two factors act in the same direction it is termed cogradient variation (positive covariance; Fig. 1d) [2,13]. Thus, when comparing among wild populations, phenotypic traits appear more similar than they would if populations were compared in a common environment for countergradient variation, and vice versa for cogradient variation.

**Figure 1:**
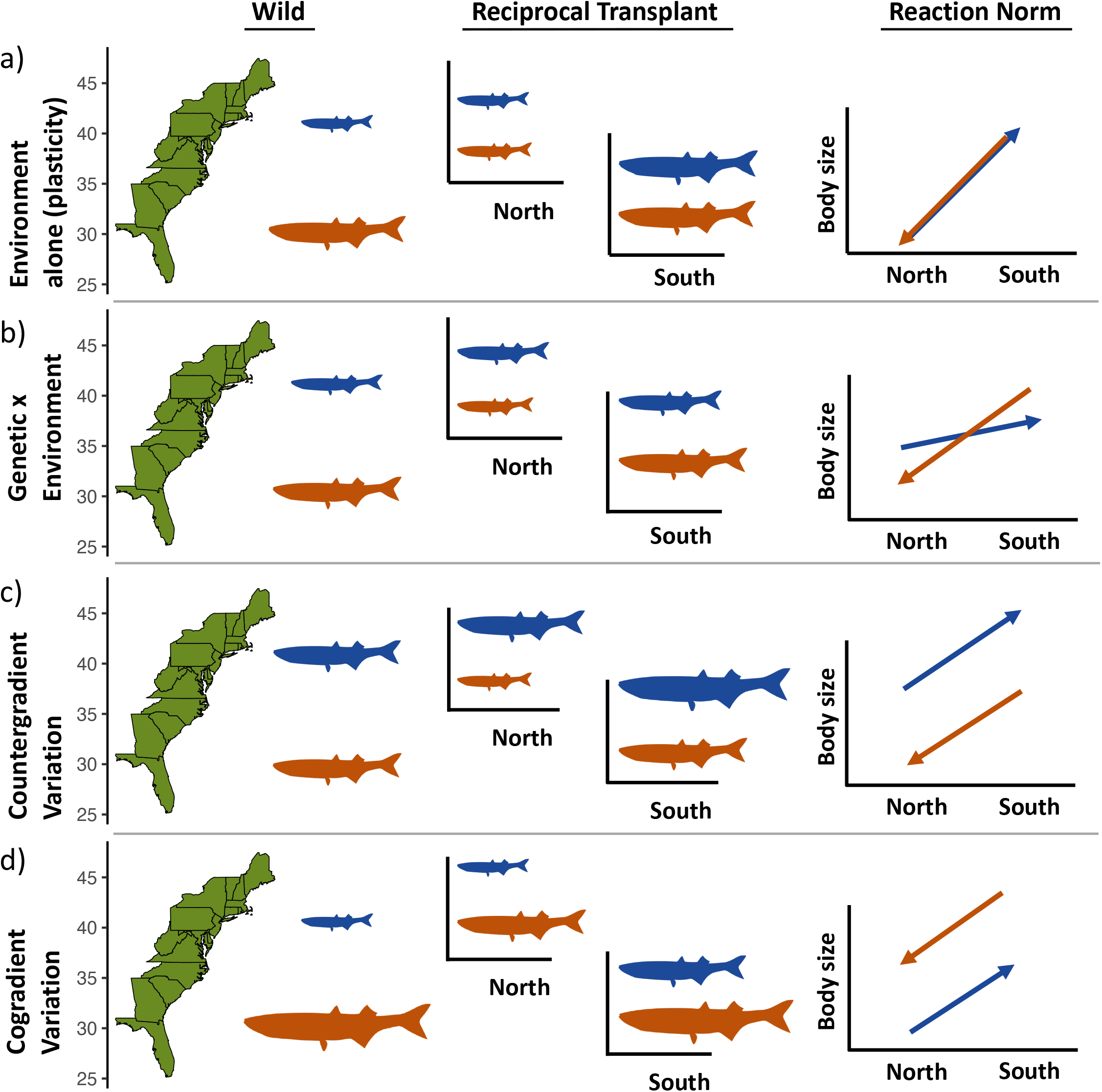
Conceptual illustration of four types of phenotypic responses for two populations of a marine fish distributed along the Atlantic coast of the United States. Each row represents four related, but different responses: a) environment alone (phenotypic plasticity without local adaptation), b) a genetic by environment response, c) countergradient variation, d) and cogradient variation. The first two columns illustrate how each of the four responses manifest in different environments—shown is the expected phenotypic responses of both populations in their home environments (“wild”) and if they were exposed to a reciprocal transplant experiment in their respective home habitats (“reciprocal transplant”). We also include the reaction norm for each response, where the base of the arrow indicates a population in its home environment and the arrowhead its away environment. Notice that for the counter-gradient variation example, both populations are the same size in the wild (*i.e*., perfect compensation), illustrating one way that local adaptation can be cryptic.

Counter- and cogradient variation constitute forms of cryptic local adaptation because their true genetic effect on phenotypic variation is masked by the environment [12–15] (Fig.1). Countergradient variation can be particularly difficult to identify in wild populations because the environmental influence on a phenotype can conceal the genetic influence, such that the signal of local adaptation only appears when individuals are reared in common environments. This response has been termed perfect compensation (or exact compensation), when wild-type phenotypes are equal across a gradient in their home environments (Fig. 1c) (*sensu* Conover *et al*. [12]). Compensation can be thought of as a measure of how similar phenotypic values are across populations in their home environments—and as an indicator of whether selection may favor similar or disparate phenotypes along an environmental gradient.

While Levins [10] was the first to formally describe countergradient variation—which was followed by early research by Berven and others [16,17] in *Ranid* frogs—it wasn’t until two-and-a-half decades later that interest in environmentally covarying local adaptation grew substantially. Research with an eastern North American fish, Atlantic silverside (*Menidia menidia*), revealed that populations had similar body sizes along a latitudinal gradient that spanned from the Gulf of Maine to the Floridian Atlantic Coast. Using a common garden approach, the researchers found that the northern populations were consistently larger than southern populations, regardless of the temperature in which they were raised (*e.g*., Fig. 1c). This finding led to substantial follow up research investigating the proximate and ultimate factors driving the trend [18,19], namely faster growth and more efficient metabolism in response to size-dependent winter mortality. A theoretical review by the group outlined the evolutionary significance of countergradient variation and described 20 prior studies with results consistent with countergradient variation, though generally without the original authors formally describing their observations as such [13]. The early work highlighted in that review and subsequent research substantiate a large body of evidence for co- and countergradient variation across diverse taxa including plants [20], bivalves [21], fishes [22-25], and insects [26-28], and across ecological gradients such as temperature [29,30], salinity [31], carotenoid availability [32,33], and urbanization [34].

Despite the attention paid to counter- and cogradient variation, there have been few studies that describe the magnitude and extent of these types of local adaptation, nor has there been much investigation into the broader abiotic and biotic factors that might contribute to their occurrence [12,35,36]. Without understanding the extent of environmentally covarying local adaptation, it is difficult to make predictions about how locally adapted populations may interact with a changing environment or how cryptic local adaptation may aid or hinder conservation goals. Here we use a Bayesian meta-analytical framework with hierarchical models to describe the general effect size of local adaptation resulting from counter- and cogradient variation as they appear in the literature and to investigate what biotic and abiotic factors are associated with the magnitude of adaptation. The effect sizes we calculate represent a direct estimate of the standard deviation of the mean difference in trait values for the local populations being measured–or more generally, they are the genetic effect of environmentally covarying local adaptation. With respect to the classic representation of phenotypic variance [37]

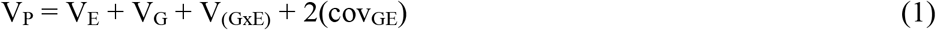

this study estimates V_G_ or the total phenotypic variance driven by genetic differences in populations that exhibit environmentally covarying local adaptation (*e.g*., cov_GE_ < 0;countergradient variation, or cov_GE_ > 0;cogradient variation, Fig. 1c,d; Fig. S1).

Counter- and cogradient variation as phenomena, represent the negatively or positively covarying contributions of V_E_ and V_G_ in equation 1 (cov_GE_). To directly measure this covariation studies are limited to reciprocal transplants or common gardens that recreate home environments [36,38]. Additionally, while cov_GE_ may be large, little research has explored how the different phenotypic variance components in equation 1 are structured along gradients in wild populations [38]. While V_G_ represents both the full effects of genetic change on phenotypic variance (including drift and gene flow), clinally covarying V_G_ of fitness related traits covary specifically because of selection and the measure of V_G_ in this specific scenario is a suitable representation of the magnitude of local adaptation in these populations. As such, the goals of our analyses were to *i*) quantify the genetic variance component of countergradient and cogradient variation as an effect size, *ii*) describe these effect sizes within and across taxa, phenotypic traits, and different gradient types, and *iii*) quantify the effect size of compensation—or how similar countergradient populations are in their home environments—to assess if there are clear trends for phenotypic similarity or dissimilarity along gradients.

## Methods

### Literature Review and Selection

Overall, we collected 858 studies that investigated counter- and/or cogradient variation. To generate a full list of studies, we first analyzed studies indicated as showing counter- or cogradient variation in the qualitative analysis from Conover *et al*. [12]. We then searched the Web of Science topic field using the terms “counter*gradient variation” in May of 2018 and “co*gradient variation” (while excluding the previous search, as the wildcard renders them redundant) in June of 2019, resulting in 384 and 34 results, respectively. In August 2019, we also included studies citing the Conover *et al*. [12] review, as well as the earlier review article by Conover & Schultz [13]—682 studies total (some of which were redundant with studies in the prior search resulting in a total of 858 studies). These methods are discussed in more depth in the Supplementary Methods.

To be further included in our analysis, a study had to meet a set of qualifying criteria. Specifically, a study had to be designed to include some form of a common environment—this may have been in a common garden study or a reciprocal transplant using wild populations or populations recently brought into lab conditions. We did not include examples from domestic populations or lab induced selection. Additionally, studies must have included two or more common environments to determine the signal of local adaptation (countergradient, cogradient, or others such as GxE that were ultimately not included in this study), as well as two or more populations to compare the genotypic difference of the phenotypic response between populations. Finally, studies were determined to show counter- or cogradient based on plotted reaction norms to verify environmental and genetic effects covaried—*i.e*., populations in away environments minimized (countergradient) or maximized genetic variance (cogradient) (positive or negative covariance, Fig. 1c,d)—and genotypes maintained their rank order across environments (Fig. S1). Occasionally, studies investigated these distinctions across multiple species in the place of populations and such studies were discarded.

### Statistical Methods

To calculate effect sizes, we took data from both tables and figures in manuscripts. For data in the form of figures, we used the software WebPlotDigitizer [39] to extract the relevant data. Briefly, the program works by importing a photo of the respective figure and the user manually sets the scales for the x- and y-axes based on the figure values and then adds digital points with their cursor to the data of interest. The digital points are then translated to numeric data via the imaging algorithm.

The statistic used to calculate effect size was Hedge’s *d*, which, for this study, was a standardized mean difference with a correction for small sample sizes[40,41] to avoid bias in the test statistic [42,43]. We calculated standardized mean difference (synonymous with effect size in this paper) by comparing the most disparate populations at each treatment level. For example, if we had three populations collected from an elevational gradient at 500m, 1000m, and 1500m in common gardens with three temperature treatments (*e.g*., 10, 15, and 20 °C, as was often the case where latitude or elevation were proxies for temperature clines), then we would collect the mean and variance for every population in every treatment, but calculate the effect size (standardized mean difference) by using the values from the 500m and 1500m populations at each treatment. Occasionally, not all populations were used in all treatments, in these scenarios we used only the treatments for the most extremely distributed (with respect to the gradient— *e.g*., very highest and lowest elevations) populations. Some studies conducted multiple experiments measuring the same trait. If multiple experiments measuring the same trait were recorded, we calculated the effect sizes for each.

In all, we calculated 422 total effect sizes from 1204 individual data points extracted from 83 studies. Many studies showed counter- or cogradient patterns in multiple traits and/or trials within a given study. We collected all these data and controlled for their non-independence using a robust, Bayesian hierarchical modeling approach (discussed below). Hedge’s *d* was calculated using the *metafor* package in R (version 3.6.3) [44] with the *escalcl()* function [45]. Additional metadata was collected from each study and included features such as year the study was published, the species investigated, and the distance in meters (for elevational gradients) and kilometers (for latitudinal gradients). The full list of these covariates is available in our supplemental metadata (Supplementary Data) and were collected to be used as covariates in the mixed-effects model (Table 1).

**Table 1:**
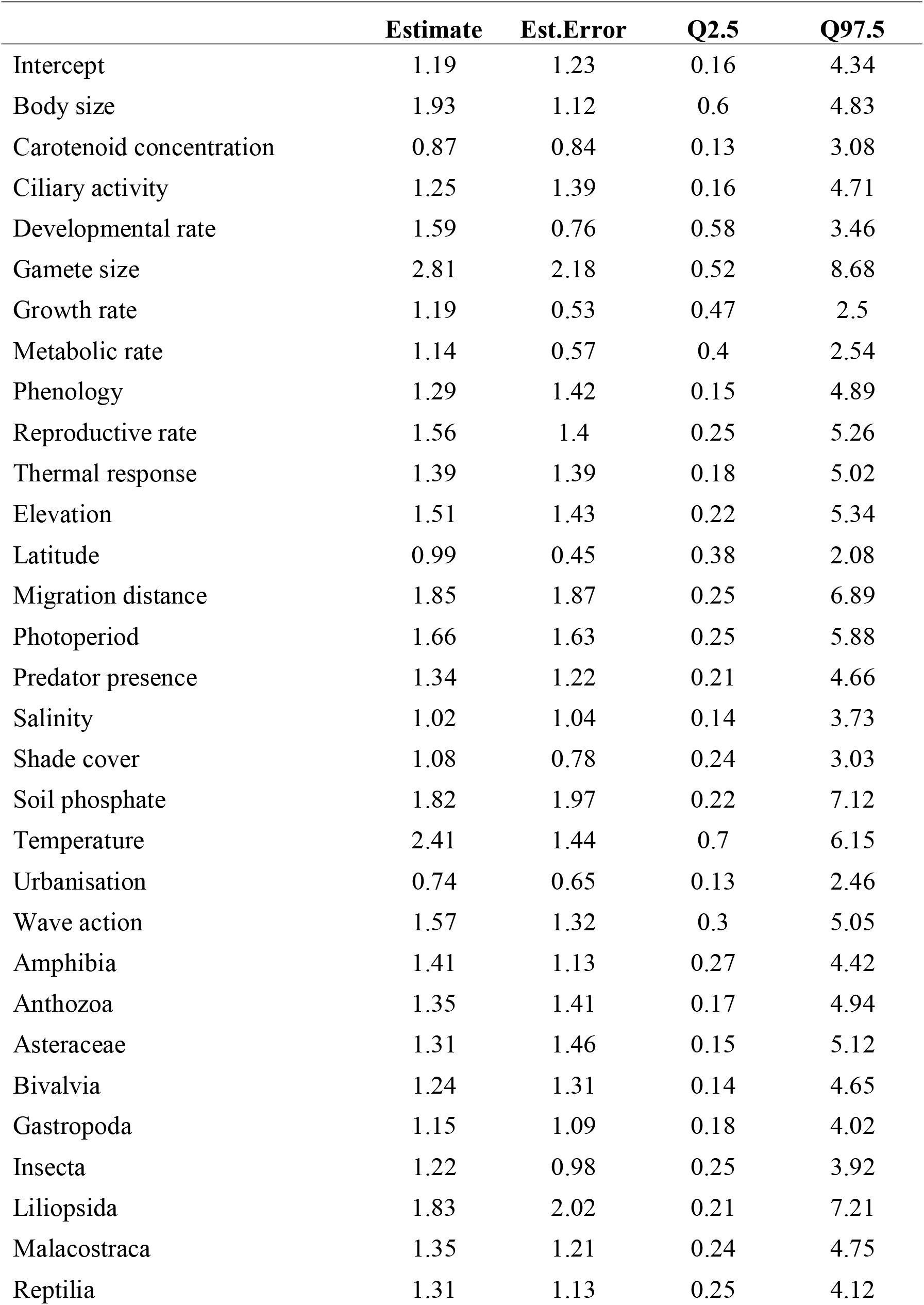
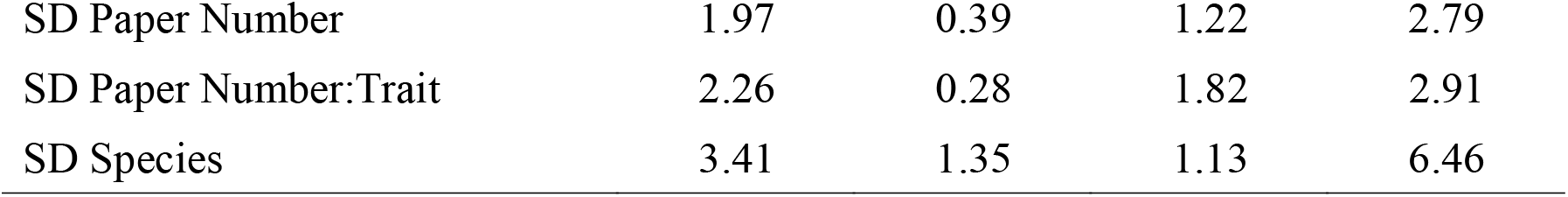
Results of Bayesian metaregression mixed model with effect size fit as a function of trait, gradient, and organism class. Random effects included trait nested in paper and paper alone, as well as a phylogenetic species effects. Shown is the intercept and the effect of each level of a covariate along with their estimated error and 95% credible intervals.

To estimate overall effect size and the effect of covariates, Bayesian linear mixed models fit with random intercepts were used in the R package *brms* [46,47], which is a high-level interface in R for the Stan modeling language [48]. Linear mixed-effects models were chosen due to the hierarchical nature of this data set, where a study may have effect sizes estimated for multiple traits in multiple experiments. For this approach, we fit the multiple estimates of effect size for a trait in an experiment by using an estimate from each experimental level (Fig. S1c)— for instance, using the prior example, an estimate for each of 10, 15, and 20 °C. This approach also allows each treatment to have different effect sizes and accounts for variability driven by Genotype x Environment interactions (V(_GxE_) eq.1). As such, nested random intercepts effect was used to control for this non-independence. We originally set out to nest traits in experiments in studies in our random effects but model parameterization necessary to overcome divergent transitions (specifically, max tree depth in Stan) extended model run time to be computationally intractable. Random effects were instead reduced to traits nested in studies.

Similarly, phylogenetic relationships may provide another source of non-independence, and was controlled for by using a phylogenetic random intercepts effect for which we created a variance-covariance matrix [49]. To do so, we generated a pseudo-phylogeny using the *taxize* package in R [50]. Using the *classification()* function, we extracted the full taxonomy of a given organism in our dataset from the NCBI database. We then used the *class2tree()* function to generate a phylogenetically accurate tree in terms of taxonomic rank (*e.g*., all species within a genus would have the same distance from one another, all genera within a family would have the same distance—a full explanation of the method is available in the Supplementary Methods). The function not only clusters taxa on their specific taxonomic rank but also within the taxonomic clade, allowing for more accurate distance measures. The tree created using this method was transformed into a matrix and then used as a custom variance-covariance component in our hierarchical model. The mathematical representation of our random effects in all models is shown in our Supplementary Materials (Supplementary Methods). We used moderately informative priors for our hierarchical models [51]—except for the metaregression, where a stronger prior was used to rein in variance—lists of priors and discussion of their role in controlling for potential bias are available in Supplementary Methods (Tables S1-S4, diagnostic plots Figures S2-5). Because we used a Bayesian framework, the following effects are presented with the means and 95 percent credible intervals (indicated as 95% CI) of the posterior distributions, unless otherwise noted.

The overall effect size for both co- and countergradient variation was estimated using random-effects only models, which is the convention when assuming study effects may vary in meta-analytical approaches [52]. We included the covariates of trait, gradient, and taxonomic class in our metaregression countergradient model to assess how different biotic and abiotic factors may contribute to the estimate of countergradient variation genetic variance effect size.

### Compensation Analysis

Finally, we analyzed whether populations appeared to be under-, over-, or perfectly compensating (*sensu* Conover *et al*. [12]) when comparing their home environment phenotypes. To do so, we used the most extreme treatments and paired them with the polarly (as in opposites, not necessarily geographically) distributed populations used in the study. For example, if we consider three populations of fish from a range of saline conditions—freshwater, brackish, and marine—then, the most saline treatment in the experiment would be considered the home environment of the marine population and the least saline the home environment of the freshwater fish. We then compared the trait values of the two polarly distributed populations and, using the previous example, if the trait value was larger for the most saline population than that of the least saline population the relationship was classified as overcompensating, close to even was classified as perfectly compensating, and negative was classified as undercompensating. To standardize these data, we again used Hedge’s *d*, because the values of different traits are generally measured on different scales [53]. Because there are no quantitative definitions of compensation, our results were conservatively classified as overcompensating if values were > 0.5, perfectly compensating if values were between −0.5 and 0.5, and undercompensating if values were < −0.5. Because compensation measurements were computed as Hedge’s *d*, these were meant to reflect a broad distribution around 0 for perfect compensation (what amounts to a medium effect size in both positive and negative directions, [54]) that encapsulates the inherent measurement error in the value.

## Results

### Random Effects Models

The result of our random effects models that estimated overall effect size for countergradient variation was 1.05 (95% CI = 0.30-2.49) and for cogradient variation 2.13 (95% CI = 0.35-7.00) (Fig. 2, non-truncated in Fig. S6). The heterogeneity parameter τ—the full random effect standard deviation—was modeled with its own prior distribution and resulted in 2.19 (95% CI = 1.79-2.82) for countergradient variation and 9.33 (95% CI = 4.00-22.45) for cogradient variation. We similarly included a random effect for phylogenetic relatedness which resulted in an estimate of 2.54 SD (95% CI = 1.04-5.28) for countergradient variation and 3.71 SD (95% CI = 1.03-14.82) for cogradient variation (Tables S5, S6).

**Figure 2:**
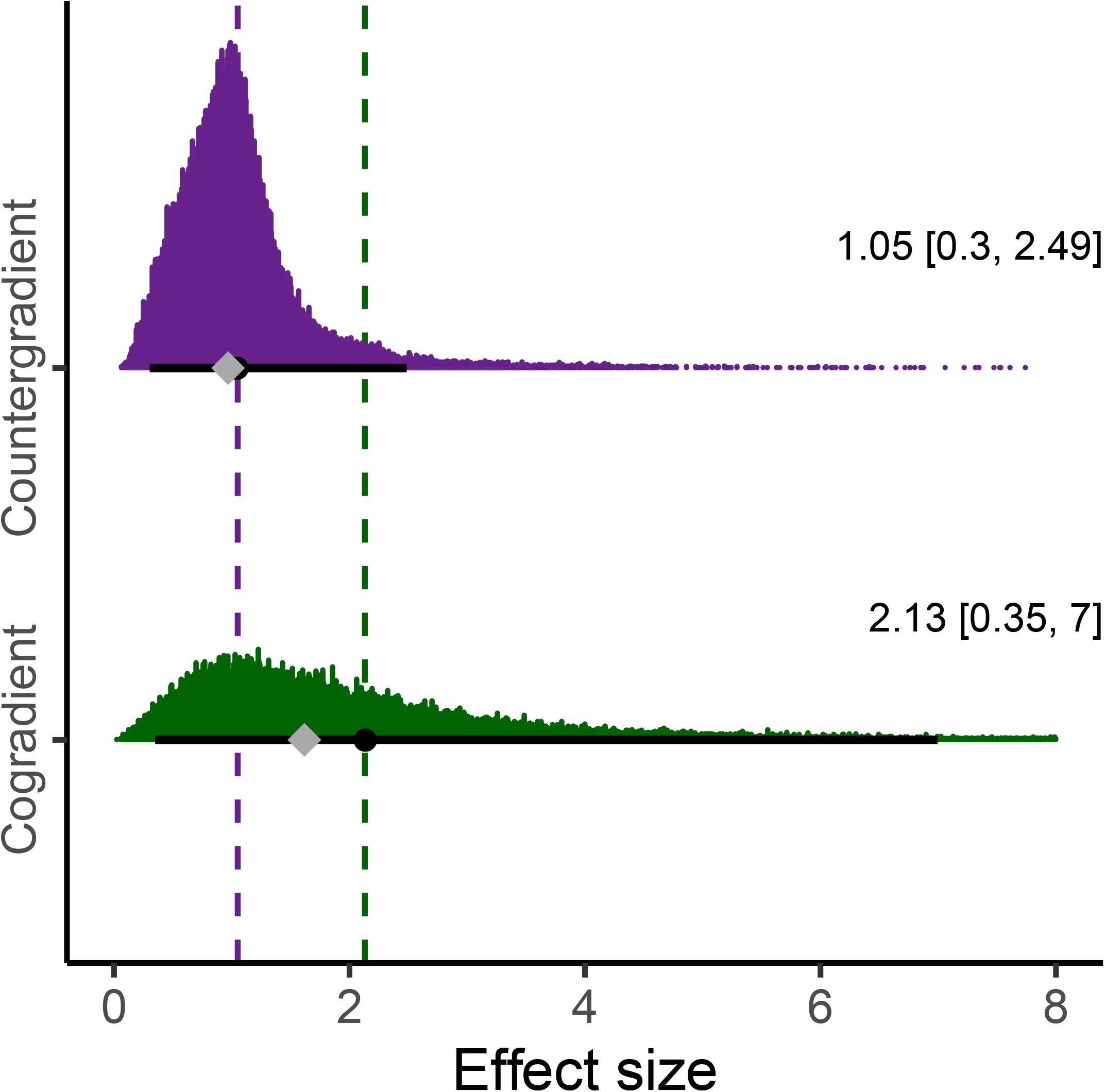
Posterior distributions of effect size estimates from random-effects meta-analysis for counter- and cogradient variation. The circle indicates the mean and the diamond the median of the posterior distribution (overlapping in countergradient) and the solid, black line, the 95% credible intervals. The mean and credible interval values are also indicated with text on the right. A total of 557 of the largest values (of 50,000 in each distribution) were removed because they extended beyond the x-axis margin (see Fig. S6 for all data points). Purple represents the posterior distribution for the effect size of countergradient variation and green, cogradient variation. The dashed line illustrates the mean value of the posterior distribution for countergradient (purple) and cogradient (green) variation.

### Metaregression

We also implemented a mixed effects metaregression with our countergradient dataset to estimate the effect size for trait classification, gradient classification, and taxonomic class effect. The mean effect, posterior distribution and 95% credible intervals for each modeled covariate are presented in Table 1 and those same values are represented as the main effects estimated marginal means with the posterior distribution and credible intervals in Figure 3 (non-truncated in Fig. S7). For each covariate, these results illustrate similar mean values with large and overlapping credible intervals. While variability was large across all covariates, the mean coefficient estimates of the posterior distributions for temperature (mean = 2.41, 95% CI = 0.70-6.15) and gamete size (mean = 2.81, 95% CI = 0.52-8.68) were much larger than the global estimate of 1.19. By contrast, the estimate for urbanization (mean = 0.74, 95% CI = 0.13-2.46)— had lower credible intervals near zero and was much lower than other estimates. These values are represented as main effect marginal means in Figure 3. The among study standard deviation was 1.97 (95% CI = 1.22-2.79), while the among trait (nested in study) standard deviation was 2.26 (95% CI = 1.82-2.91), and the standard deviation for phylogenetic random effect was 3.41 (95% CI = 1.13-6.46), indicating more variance was accounted for among traits in studies and by phylogeny, than among studies.

**Figure 3:**
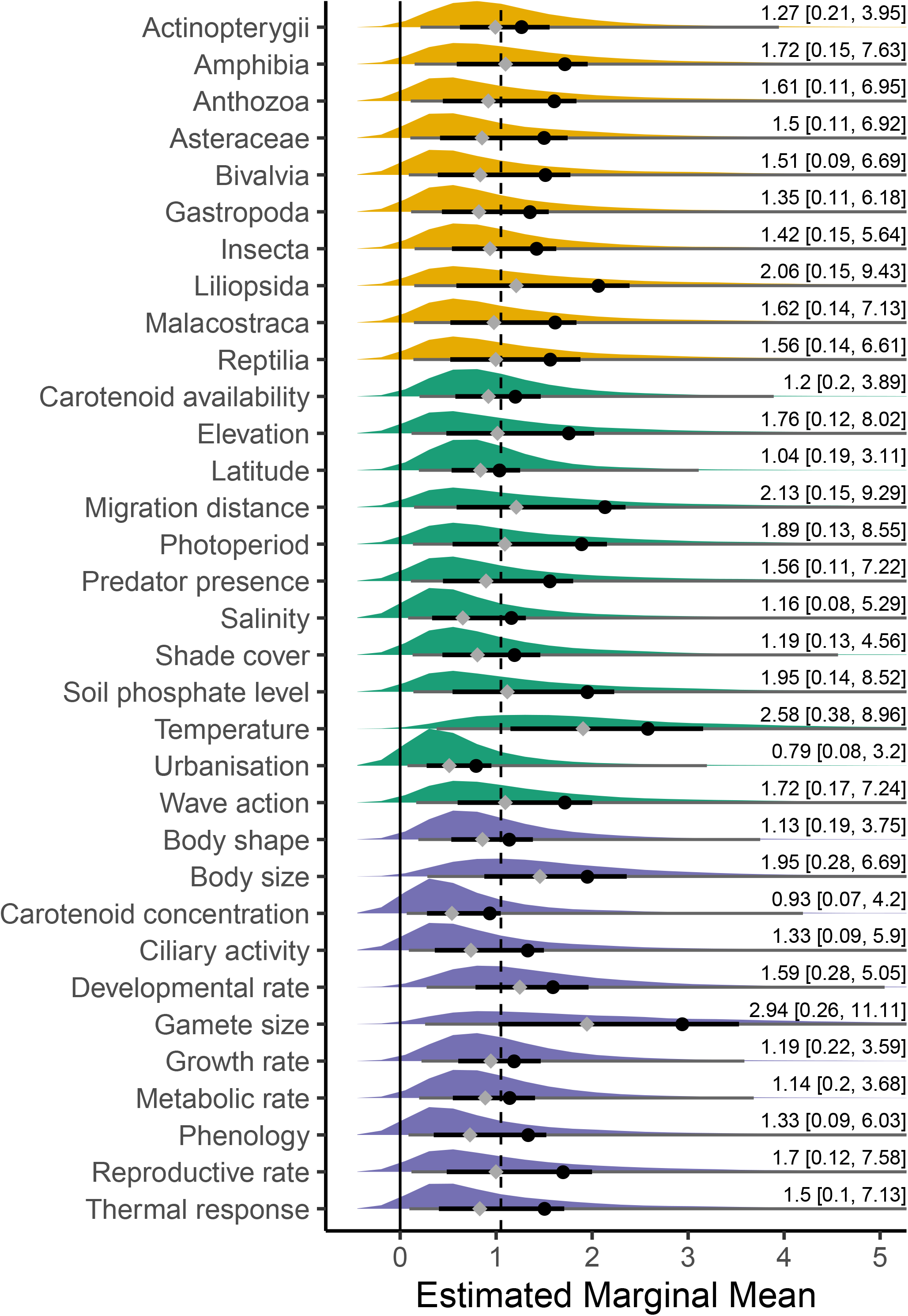
Posterior distributions of estimated marginal means for the covariates in a metaregression analysis for countergradient variation. Yellow distributions depict taxonomic class, green distributions depict environmental gradients, and purple distributions depict trait covariates. The black circle indicates the mean and grey diamond the median of the posterior distribution. The thick solid, horizontal black line depicts the 50% credible intervals for each covariate and the grey line the 95% credible intervals, some of which extend beyond the x-axis limit (the full 95% CI is indicated with text to the right). The x-axis limit was set to 5 to better visualize the behavior around the bulk of the posterior, as such 8,949 of the largest values (of 60,000 thinned to 6,000 in each distribution—198,000 total) were removed because they extended beyond the x-axis margin (all values can be viewed in Figure S7). The dashed line illustrates each posterior distribution’s relationship to the study-wide effect size of 1.05 estimated from the random effects model and shown in Fig. 2.

### Compensation Analysis

The results of our compensation analysis were plotted to represent the conceptual relationship between home phenotypes of populations exhibiting countergradient variation (*sensu* Conover *et al*. [12]) (Fig. 4). Our data provide support for far more frequent over- and under compensation than perfect compensation, meaning that populations in our study generally have divergent home phenotypes as opposed to identical phenotypes along a gradient. We considered any value between −0.5 and 0.5 as perfectly compensating (or a range of 1 effect size around 0 for a conservative estimate), values less than −0.5 undercompensating, and values larger than 0.5 overcompensating. We consider this classification conservative because it allows for a medium effect size in the positive or negative direction from zero [52], despite the theoretical description of perfect compensation being a value equal to zero [12]. The distribution of these values in our data set were overcompensating = 38.6%, undercompensating = 36.0%, and perfectly compensating = 25.4%. Effect sizes of compensation ranged from −70.38 to 36.45 but 5% and 95% quantiles were −27.74 and 6.49, respectively.

**Figure 4:**
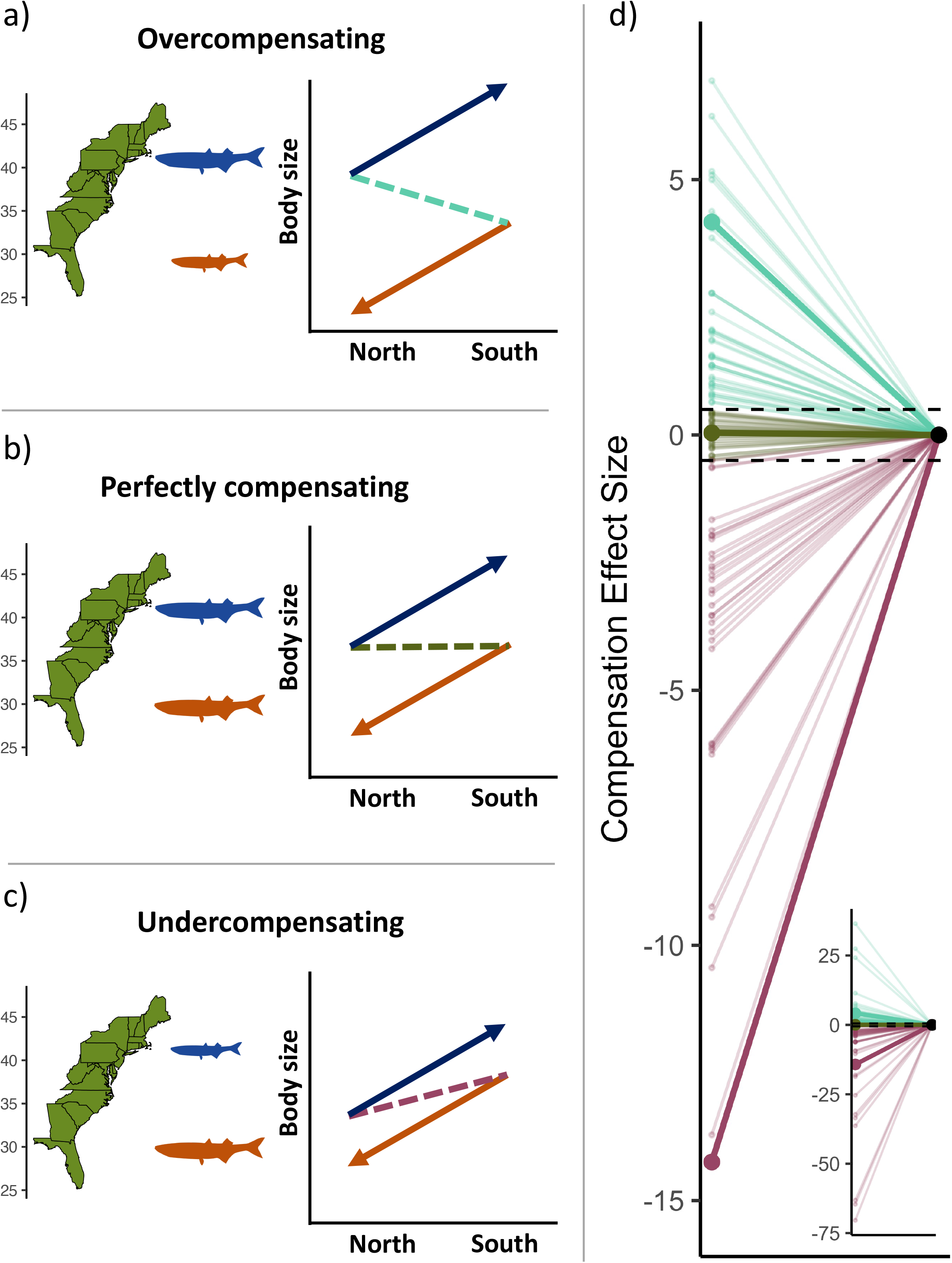
Conceptual representation of two countergradient populations in their home environments and the reaction norms of those populations in the home and away environments. Also shown are the results of our compensation analysis. a) Represents an overcompensating relationship where the northern population is larger than the southern population in all environments and the slope of the dashed line connecting their home phenotype (arrow bases) has a negative sign as it decreases from the larger northern population to the smaller southern population. b) Represents a perfectly compensating relationship, where phenotypes are equal in their respective home environments and thus the slope is flat. c.) Represents an undercompensating relationship, where the southern population is larger than the northern population in their respective home environments and the slope of the dashed line connecting their home phenotype (arrow bases) has a positive sign as it increases from the smaller northern population to the larger southern population. d.) represents the results of our compensation analysis where overcompensating was defined as an effect size > 0.5, perfect as 0.5 ≤ and > −0.5, and undercompensating < −0.5. Of these three responses, our data was distributed as 38.6% overcompensating, 25.4% perfectly compensating, and 36.0% undercompensating. Points on the left are the calculated effect sizes and are connected to a point at 0 to show a slope to match the conceptual panels. The larger figure is bounded at −15 and 7 (removing 7 values) to better show the results near 0 and perfect compensation, whereas the inset figure includes all data.

## Discussion

The ecological and evolutionary importance of co- and countergradient variation has been recognized for three decades [13,18]. Here, we find that the genetic effect of populations expressing counter- and cogradient variation are 1.05 and 2.13, respectively. Our measure of standardized mean difference, Hedges’ *d*, translates the difference between populations into units of standard deviation. Thus, our results indicate that the standardized mean difference between the most disparate populations in our study—*i.e*., the genetic effect of environmentally covarying local adaptation—is between one and two standard deviations change in trait value. In other words, populations that express the same trait value in their home environments would diverge by one or two standard deviations in a common environment due to their genetic differences depending on whether they were expressing counter- or cogradient variation, respectively. Additionally, our results suggest populations expressing cogradient variation may express twice the genetic divergence as those expressing countergradient variation, but the results for cogradient variation are based on small sample set with significant variance. Our findings demonstrate that the genetic divergence among locally adapted, clinally distributed populations is large but also obscured as a function of the covariance between environmental and genetic effects and their interaction with phenotype.

We used a metaregression to investigate the effect sizes of abiotic and biotic covariates of countergradient variation. Covariates were statistically indistinguishable from one another due to high variance in their estimates, though many showed mean values that were large with distributions away from 0 (Effect sizes estimates of 1-3, Table 1, Fig. 3). Particularly, we estimated large effect sizes for gamete size (2.81), and temperature (2.41). For gamete size, multiple studies in fishes showed effects greater than 2.5 and as high as 13.41 [55]. The tradeoff between size and number of eggs produced by female fishes is a well-established trend in ecology [56,57], and particularly in marine fishes where there are strong links between egg size and latitude [58]. The large effect size estimate for gamete size from our metaregression then is less surprising given the strong selection on gamete size across species and environments, but also suggests that despite the tradeoff between size and number of eggs there is benefit to larger eggs moving poleward. While there were additional large study-specific effects for temperature (4 instances > 4.5), we separated temperature and temperature related gradients based on the explicit distinction made by authors. Temperature is a fundamental selective agent for wild organisms, while latitudinal, elevational, and shade cover gradients serve as proxies for temperature—all three had estimates at or near one standardized mean difference. The proxy measures then likely fail to capture the full evolutionary dynamics driving environmentally covarying local adaptation at the same resolution as that caused by temperature.

As far as the variability in our metaregression estimates, we find that the inter-effect variability (τ^2^, or the variability of our random effects) generally swamped the covariate-specific effects estimates (Table 1). The variability in this random effect is also a measure of GxE (the variability of multiple effect sizes for a trait in a study, Fig. Sc) and indicates that when environmentally covarying local adaptation is present, this adaptation corresponds with a large population-specific (or genotype-specific) GxE influence. While the lack of predictive results in our metaregression may be indicative that our data is not robust enough for differences among covariates to emerge, we think it is more likely that these results suggest that countergradient variation, when it does occur, is an ecologically ubiquitous phenomenon for species distributed along environmental gradients. Furthermore, we suspect that species or populations experiencing similar agents of selection—for example, those along latitudinal, elevational, or temperature gradients—may achieve adaptation through different mechanisms (as indicated by the large GxE influence) and therefore contributing to the variability observed in our models. In other words, while the phenotypic solution may appear similar across species and gradients, the processes driving the genetic divergence of the phenotypes of different populations is idiosyncratic. The conclusion that covariate specific effects are idiosyncratic is further supported by the lack of pattern with respect to latitudinal distance and temperature difference regressions analyses, neither of which were predictive (Fig. S8, Table S7). Thus, while populations almost always showed a large effect of environmentally covarying local adaptation—metaregression estimates were 2.41 and 0.99 for temperature and latitudinal gradients, respectively—the magnitude did not scale with temperature or physical distance based on separate analyses (Fig. S8, Table S7). This may be due in part to microgeographic variation—in which large adaptive phenotypic divergence occurs over small spatial scales—as multiple studies reviewed as part of this work indicated microgeographic variation as driver of phenotypic patterns across the landscape [59,60].

Even though the overall effect size of cogradient variation was large, only 15 out of 858 studies were consistent with cogradient variation and included enough information to calculate effect sizes. This paucity may reflect an overall lack of published research investigating and demonstrating cogradient variation (see also, [12]). However, in a meta-analysis of reciprocal transplant studies, Stamp & Hadfield [36] found 60% of traits exhibited a cogradient pattern. Our results (*i.e*., the magnitude of effect size), in conjunction with the work of Stamp & Hadfield [36] suggest that, while rarely reported, cogradient variation constitutes an ecologically meaningful phenomenon. We further hypothesize cogradient variation may suffer from underreporting because it’s natural sign mirrors environmentally induced plasticity that may not drive researchers to investigate populations using common gardens. While cogradient variation may be underreported, standalone cases such as that by Trussell & Etter [61] can show both the large effect of cogradient variation and how home phenotypic differences can disguise that effect. In a reciprocal transplant study, the researchers measured shell thickness in marine snail populations, finding that, when in their home environments, populations expressed a 3.18 standardized mean difference in shell thickness. But, when considered in shared environments, that difference reduced to 2.07 units. Notably, this result amounts to a substantial effect size difference due to genetic variation alone, but is ~2/3 value if the populations had only been observed solely in their home environments. More importantly, if the researchers had only investigated these populations in their home environments without common gardens, a likely inference for the phenotypes would have been a strong plastic response, which, while true, did not account for even half of phenotypic variation in home environments.

Our compensation analysis for countergradient variation revealed that populations in their home environments were more commonly over- or undercompensasting, as opposed to perfectly compensating (Fig. 4). In other words, the genetic effect of countergradient variation was generally not equal to that of the environmental effect. This result was clear with conservative bounds for perfect compensation (−0.5 to 0.5 standardized mean difference). Reducing those bounds to the less conservative values of −0.2 to 0.2 (considered to be a small effect size [54]) for perfect compensation resulted in only 11.7 percent of studies demonstrating perfect compensation. While we caution that these results may not necessarily point to clear biological trends about how common and in what scenarios over- and under-compensation may occur, it does suggest that either: 1. selection for maintaining optimal phenotypes across broad environmental gradients may not be the norm *(sensu* [12,13]), or 2. the response to selection may not be strong enough to effectively minimize phenotypic variation across populations. If the latter, the response to selection may be minimized by insufficient adaptive genetic variation or countervailing selection on correlated traits specific to each local population [62].

Our results are indicative of a response to selection contributing to strong genetic divergence in trait values distributed along environmental clines and reinforce the evolutionary importance of counter and co-gradient variation. As such, negative or positively environmentally covarying selection is a strong candidate as a mechanism for driving parapatric speciation—if the selection pressure is consistent across the gradient—or peripatric speciation—if selection is strongest at the poles. Genetic differentiation may also be expected by further anthropogenic change, where habitat fragmentation breaks up corridors of gene flow within clines [63], or climate change disrupts habitat envelopes at the polar ends. Both scenarios have significant implications for how populations exhibiting environmentally covarying local adaptation are managed. This clinal breakdown may also lead to underestimation of the importance of co- and countergradient variation, as range-wide patterns may have already been disrupted causing researchers to undercount their occurrence or underestimate their historical magnitude if the edges of clines are lost. Additionally, environmentally covarying adaptation should be considered in conservation and management efforts, as cryptic local adaptation may lead to undesirable management outcomes if ignored, or good candidates for translocation (*e.g*., genetic rescue [64]) if well understood.

## Conclusion

Our findings illustrate that environmentally covarying local adaptation is pervasive across multiple gradients in a diverse set of taxa. With this meta-analysis, we quantitatively characterized the genetic effect on phenotypic divergence for environmentally covarying local adaptation and find that the result is one to two full standard deviations from the mean trait value of a population. While the genetic effect of co- and countergradient variation may be large, the variability of response observed across studies is also quite high, as indicated in the values of our random effects, indicating strong GxE influences. Moreover, when analyzing the mean trait difference between home phenotypes of populations exhibiting countergradient variation, we find no support for a strong signal of perfect compensation, indicating that selection for optimal phenotypes across a gradient can or does not perfectly minimize variation in mean phenotypic trait values. Much like local adaptation more generally, environmentally covarying local adaptation necessitates close consideration when working with populations distributed across ecological gradients. We conclude that environmentally covarying local adaptation is ubiquitous with a large effect size and argue that this form of cryptic local adaptation deserves continued attention with theoretical, experimental, and genomic approaches.

## Supporting information

Supplemental Metadata Table

## Acknowledgements

We thank K. Lotterhos, N. Emery, and C. Oakley for providing helpful discussion regarding environmentally covarying local adaptation. K. Horvath offered useful suggestions about instituting a phylogenetic random effect in our models, D. Pascall provided valuable insights on implementing the pseudo-phylogeny, and R. Swihart gave useful direction on Bayesian model setup. We also thank A. Harder, A. Lee, and A. Nalesnik for their helpful comments on a past version of this manuscript. This work was supported, in part, by the Purdue Ecology and Evolutionary Biology Waser Fellowship to MMS and by funding from the National Science Foundation (OCE-1924505; DEB-1856710) to MRC.

