## Supplementary Materials for "Large genetic divergence underpins cryptic local adaptation across ecological and evolutionary gradients"

**Supplementary Methods:**

*Literature Review and Selection*

For this meta-analysis, we followed strict guidelines outlined by the Preferred Reporting Items for Systematic Reviews and Meta-Analyses (PRISMA) (Page *et al.* 2021). We used the 2009 version of the checklist, as data collection took place prior to the updated 2020 version, which remained similar but with more emphasis on automized search methods and pre-reporting planned methods (Page *et al.* 2021). All studies used where effects were calculated are cited below.

Furthermore, each study must have included some mean estimate for a population phenotypic value (or data that could be used to calculate a mean), a measure of variation that could be transformed into units of standard deviation, and sample sizes for these statistics. When studies reported ranges an estimate for SD was made by subtracting the maximum value from the minimum value and then dividing by four as in Davidson *et al.* (2011).

We used the absolute value of the collected effect sizes because sign was a function of how a measurement was taken. For instance, consider a countergradient cline of populations distributed over a thermal gradient. The population in the coldest environment would always display the fastest growth, hence a positive sign, but when measuring that same population for developmental rate (days to hatch), a countergradient response would be fewer days and hence a negative sign. This is true for cogradient populations, as well. Additionally, while counter- and cogradient responses may be the inverse of one another with respect to their reaction norms, we are attempting to describe the magnitude of the genotypic difference and when considering how different traits affected sign, we felt it best to express those differences as absolute values. The statistical approach with the log transformation and the use of the Student’s T-distribution are to account for using absolute values and that only certain distributions (normal, Student’s T) allowed weighting on variance in the *brms* packge.

While we did not solely include studies that explicitly measured fitness or traits experimentally linked to fitness, counter- and cogradient variation constitute a special case of local adaptation where the genetic component of phenotypic variance is structured along clines due to environmentally covarying selection. This process is the interaction between counter- or cogradient selection with counter- or gradient variation as the adaptive response outlined in Conover *et al.* (2009) and such selection and adaptation has been demonstrated in many of the prominent case studies of environmentally covarying local adaptation.

*Statistical Methods*

The model format for the random effects models for counter- and cogradient variation is expressed below, the random effect structure stayed consistent for all models throughout our study:

Effect Size_i_∼*N*(α_j[i],k[i],l[i]_,σ^2^)

α_j_∼*N*(μ_αj_,σ^2^_αj_), for Trait:Paper Number j = 1,…,J

α_k_∼*N*(μ_αk_,σ^2^_αk_), for Paper Number k = 1,…,K

α_l_∼*N*(μ_αl_,σ^2^_αl_), for Species l = 1,…,L

where *j* is trait nested in paper, *k* is paper number, and *l* is the phylogenetic random effect.

*Prior Designation and Controlling for Potential Bias*

For our models, we used four chains, with thousands of burn-ins, tens-of-thousands of iterations for each chain, with thinning set to 1 to 10 depending on the number of iterations in the model, resulting in >5,000 estimates for our posterior distributions (exact values are available in Table S4, diagnostic plots are included in Figures S1-4).

The overarching goal of this study was to estimate the genetic divergence of populations exhibiting environmentally covarying local adaptation. The cryptic nature of these phenomena may mean they are not described as commonly as they occur in nature, but also, because their occurrence is not taxonomically or ecologically limited, the moderate number of examples we have are spread across a multitude of ecological covariates. As such, there is some difficulty translating those data into statistical models. To account for those factors, we used a robust Bayesian modelling approach to better control for non-independence—a frequentist approach would have lacked power for the metaregression—but also to account for potential publication bias in our data. For example, our priors, while weakly informative were centered around zero—which we would interpret as a lack of genetic divergence and no effect. All other factors being the same, studies with small effect sizes are less likely to be described as significant by their authors and may be missed in literature searches due to a file drawer effect. But our Bayesian approach, where priors are centered on a null effect, helps account for that potential bias and especially penalizes covariates where there are few examples to generate likelihood distributions in the model. While still imperfect, we believe this approach more appropriately accounts for the potential biases inherent in ecological meta-analyses and allows us to implement methods such as the phylogenetic random effect, which would otherwise be unavailable for these data.

*Creating Study Phylogeny*

The *class2tree()* function from the taxize package works by creating a tree based on phylogenetic rank. For example, all species within a genus would be equally distant from one another and all genera within a family would also be equally distant, and so on. While this format creates an imperfect phylogeny, it does account for phylogenetic relatedness and we do not know of any phylogeny that accounts for the full taxonomic breadth of species used in this study. A full explanation by the authors of the methodology can be found at <https://github.com/ropensci/taxize/issues/849>.

*Latitudinal and Elevational Analyses*

A large portion of our data used temperature treatments (e.g., as proxies for gradients such at latitude, elevation, or as direct reflection of temperature) or reported the latitudinal or elevation points for where their populations were collected in the wild. For these studies, we regressed effect size as a function of “home” temperature differences in a study—home being the coldest and warmest temperatures with respect to the most extremely distributed populations—and latitudinal straight-line distance. We had three studies that included elevation and were therefor not able to investigate the relationship between effect size and elevation.

**Supplementary Results**

*Latitudinal and Elevational Analyses*

To investigate how countergradient responses scaled across increasing distance or temperature gradients, we regressed effect size as a function of latitudinal distance and experimental temperature difference (Fig. S8). Neither was significant at alpha = 0.05. That is, even though there was a large average effect size across environmental gradients (Fig 3), the magnitude of this effect size was not driven by the magnitude of temperature or spatial differences (Fig. S8).

**Supplementary Discussion**

One of the confounding aspects of investigating clinal trends in locally adapted populations is the often non-continuous nature of their phenotypic values. For example, the non-continuous nature of population responses is particularly clear in Hice *et al.* (2012), in which they investigated 37 sites across the entire range of Atlantic silversides and found an overall trend of countergradient variation in growth rate, but multiple breakpoints along the cline. The results presented in Hice *et al.* (2012), constitutes one of the few attempts to characterize environmentally covarying local adaptation across the full environmental gradient for a species or set of populations. However, if the full distribution of populations is ignored or nor correctly accounted for, such sample designs may present a false signal of environmentally covarying local adaptation. Few studies in this analysis included measures of genetic distance among the populations they were measuring. For example, hypothetical meta-populations distributed along a gradient such as latitude with little or no gene flow between them may experience divergent selection on the same trait but from different modes of selection and could appear as environmentally covarying local adaptation.

Clinal variation has been the subject of numerous neutral and quantitative genetic studies (Richter-Boix *et al.* 2010, Frei *et al.* 2014, Hangartner *et al.* 2012), however, the pace of genomic investigation into environmentally covarying local adaptation lags other forms of local adaptation. In fact, fundamental theory about countergradient variation often assumes directional selection drives the founding populations with low relative fitness opposite of the environmental effect to achieve the negative covariance that characterizes countergradient variation. However, recent work by Wilder *et al.* 2020 found substantial genomic divergence at specific chromosomal sites among northern and southern populations of Atlantic silverside, despite these being continuously distributed populations with substantial gene flow and low genetic differentiation. These most differentiated genomic sites were largely driven by a single population in the warmest temperatures. Despite this exciting initial work, the genomic processes that drive counter- and cogradient variation, and what tradeoffs may exist in shaping the development of environmentally covarying local adaptation remain poorly understood.

**Supplementary Tables:**

**Table S1.** Prior designations for countergradient random effects model. The distribution of the prior (prior), the coefficient class (class), the specific coefficient (coefficient), the grouping format (group), and whether the prior specification was custom set or a default setting (source) is provided.

| **Prior** | **class** | **coef** | **group** | **source** |
| --- | --- | --- | --- | --- |
| normal(0, 2) | Intercept |  |  | user |
| gamma(2, 0.1) | nu |  |  | default |
| cauchy(0, 2) | sd |  |  | user |
| cauchy(0, 2) | sd |  | paper_number | default |
| cauchy(0, 2) | sd | Intercept | paper_number | default |
| cauchy(0, 2) | sd |  | paper_number:Trait | default |
| cauchy(0, 2) | sd | Intercept | paper_number:Trait | default |
| cauchy(0, 2) | sd |  | Species | default |
| cauchy(0, 2) | sd | Intercept | Species | default |

**Table S2**. Prior designations for cogradient random effects model. The distribution of the prior (prior), the coefficient class (class), the specific coefficient (coefficient), the grouping format (group), and whether the prior specification was custom set or a default setting (source) is provided.

| **Prior** | **class** | **coef** | **group** | **source** |
| --- | --- | --- | --- | --- |
| normal(0, 2) | Intercept |  |  | user |
| gamma(2, 0.1) | nu |  |  | default |
| cauchy(0, 2) | sd |  |  | user |
| cauchy(0, 2) | sd |  | paper_number | default |
| cauchy(0, 2) | sd | Intercept | paper_number | default |
| cauchy(0, 2) | sd |  | paper_number:Trait | default |
| cauchy(0, 2) | sd | Intercept | paper_number:Trait | default |
| cauchy(0, 2) | sd |  | Species | default |
| cauchy(0, 2) | sd | Intercept | Species | default |

**Table S3.** Prior designations for countergradient metaregression model. The distribution of the prior (prior), the coefficient class (class), the specific coefficient (coefficient), the grouping format (group), and whether the prior specification was custom set or a default setting (source) is provided.

| **Prior** | **class** | **coef** | **group** | **source** |
| --- | --- | --- | --- | --- |
| normal(0, 1) | b |  |  | user |
| normal(0, 1) | b | alt_traitbody_size |  | default |
| normal(0, 1) | b | alt_traitcarotenoid_concentration |  | default |
| normal(0, 1) | b | alt_traitciliary_activity |  | default |
| normal(0, 1) | b | alt_traitdevelopmental_rate |  | default |
| normal(0, 1) | b | alt_traitgamete_size |  | default |
| normal(0, 1) | b | alt_traitgrowth_rate |  | default |
| normal(0, 1) | b | alt_traitmetabolic_rate |  | default |
| normal(0, 1) | b | alt_traitphenology |  | default |
| normal(0, 1) | b | alt_traitreproductive_rate |  | default |
| normal(0, 1) | b | alt_traitthermal_response |  | default |
| normal(0, 1) | b | ClassAmphibia |  | default |
| normal(0, 1) | b | ClassAnthozoa |  | default |
| normal(0, 1) | b | ClassAsteraceae |  | default |
| normal(0, 1) | b | ClassBivalvia |  | default |
| normal(0, 1) | b | ClassGastropoda |  | default |
| normal(0, 1) | b | ClassInsecta |  | default |
| normal(0, 1) | b | ClassLiliopsida |  | default |
| normal(0, 1) | b | ClassMalacostraca |  | default |
| normal(0, 1) | b | ClassReptilia |  | default |
| normal(0, 1) | b | Gradientelevation |  | default |
| normal(0, 1) | b | Gradientlatitude |  | default |
| normal(0, 1) | b | Gradientmigrationdistance |  | default |
| normal(0, 1) | b | Gradientphotoperiod |  | default |
| normal(0, 1) | b | Gradientpredator |  | default |
| normal(0, 1) | b | Gradientsalinity |  | default |
| normal(0, 1) | b | Gradientshadecover |  | default |
| normal(0, 1) | b | Gradientsoilphosphatelevel |  | default |
| normal(0, 1) | b | Gradienttemperature |  | default |
| normal(0, 1) | b | Gradienturbanisation |  | default |
| normal(0, 1) | b | Gradientwaveaction |  | default |
| normal(0, 1) | Intercept |  |  | default |
| gamma(2, 0.1) | nu |  |  | default |
| cauchy(0, 2) | sd |  |  | user |
| cauchy(0, 2) | sd |  | paper_number | default |
| cauchy(0, 2) | sd | Intercept | paper_number | default |
| cauchy(0, 2) | sd |  | paper_number:Trait | default |
| cauchy(0, 2) | sd | Intercept | paper_number:Trait | default |
| cauchy(0, 2) | sd |  | Species | default |
| cauchy(0, 2) | sd | Intercept | Species | default |

**Table S4.** Stan control variables to define models. Iterations is the number of draws per chain (cores), Burn.in is the number of draws before sampling, and thinning refers how often draws are sampled (1 = every value, 10 = every tenth value). Adapt_delta and max_treedepth are Stan-specific variable referring to step size for the algorithm and model run depth, respectively.

| **Model** | **Iterations** | **Burn.in** | **Cores** | **Thin** | **adapt_delta** | **max_treedepth** |
| --- | --- | --- | --- | --- | --- | --- |
| Countergradient Random Effects | 20000 | 7500 | 4 | 1 | 0.995 | 20 |
| Cogradient Random Effects | 10000 | 2500 | 4 | 1 | 0.99 | 18 |
| Countergradient Metaregression | 22500 | 7500 | 4 | 10 | 0.995 | 20 |

**Table S5.** Model coefficients for countergradient random effects model indicating the intercept (overall effect) and the random effects, as well as estimated error and 95% credible intervals.

|  | **Estimate** | **Est.Error** | **Q2.5** | **Q97.5** |
| --- | --- | --- | --- | --- |
| b_Intercept | 1.05 | 0.58 | 0.30 | 2.49 |
| sd_paper_number__Intercept | 2.16 | 0.36 | 1.47 | 2.91 |
| sd_paper_number:Trait__Intercept | 2.19 | 0.26 | 1.79 | 2.82 |
| sd_Species__Intercept | 2.54 | 1.24 | 1.04 | 5.28 |

**Table S6.** Model coefficients for cogradient random effects model indicating the intercept (overall effect) and the random effects, as well as estimated error and 95% credible intervals.

|  | **Estimate** | **Est.Error** | **Q2.5** | **Q97.5** |
| --- | --- | --- | --- | --- |
| b_Intercept | 2.13 | 2.39 | 0.35 | 7.00 |
| sd_paper_number__Intercept | 3.67 | 4.87 | 1.04 | 14.10 |
| sd_paper_number:Trait__Intercept | 9.33 | 5.78 | 4.00 | 22.45 |
| sd_Species__Intercept | 3.71 | 16.96 | 1.03 | 14.82 |

**Table S7:** Model results (coefficient estimate, confidence interval, and p-value) for frequentist mixed models with latitudinal distance or experimental temperature change as predictors of effect size. Neither latitudinal distance, nor experimental temperature were significant predictors of change in effect size.

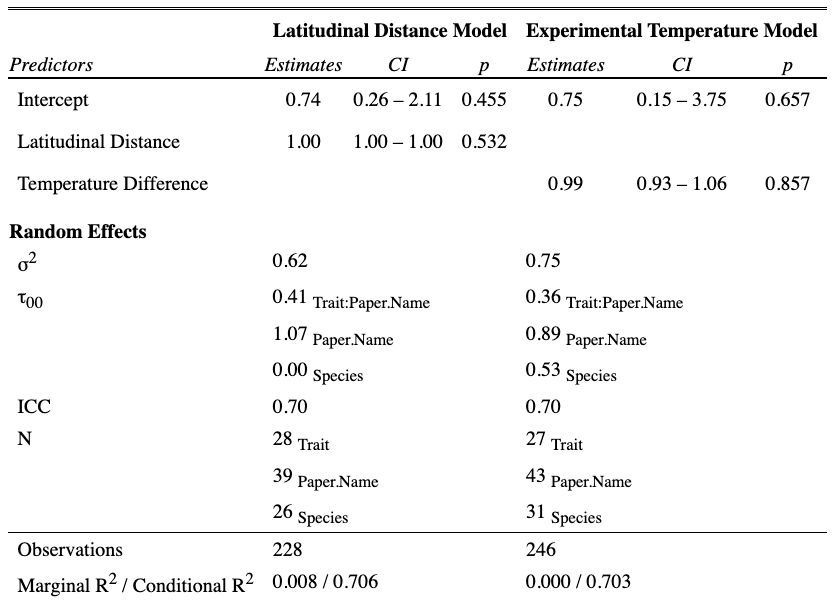

**Supplementary Figures**

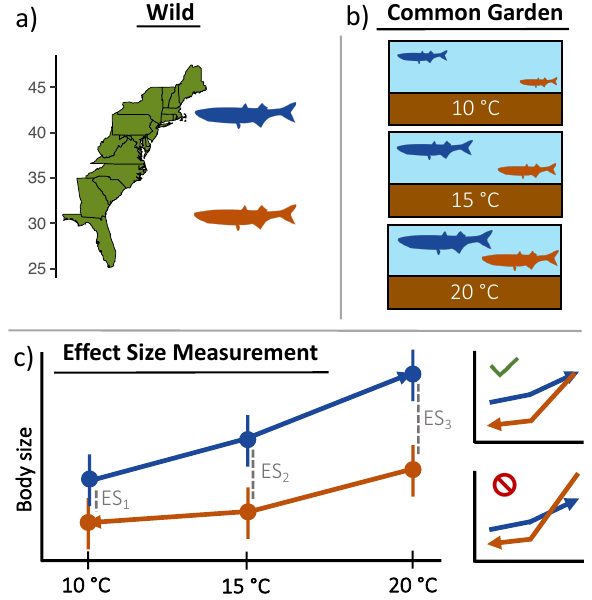

**Figure S1.** Schematic representation of data collection and analysis for meta-analysis. Panel a) represents how populations would appear in their home environments in the wild, panel b) represents a hypothetical common garden experiment researchers would have designed for their original study, and panel c) represents how the data collected in that study would have been portrayed and the effect sizes we would have calculated from that data—e.g., the mean difference at each experimental level while accounting for the variance (ES = Hedge’s *d*), note that each ES estimate could vary due to GxE. These three effect sizes would then be fit into our Bayesian hierarchical models to estimate the effect for the specific trait nested in study. The insets in panel c) represent different presentations of reaction norms and how rank order must be maintained to be interpreted as counter- or cogradient variation (green check = accepted, red crossed circle = rejected) in this study.

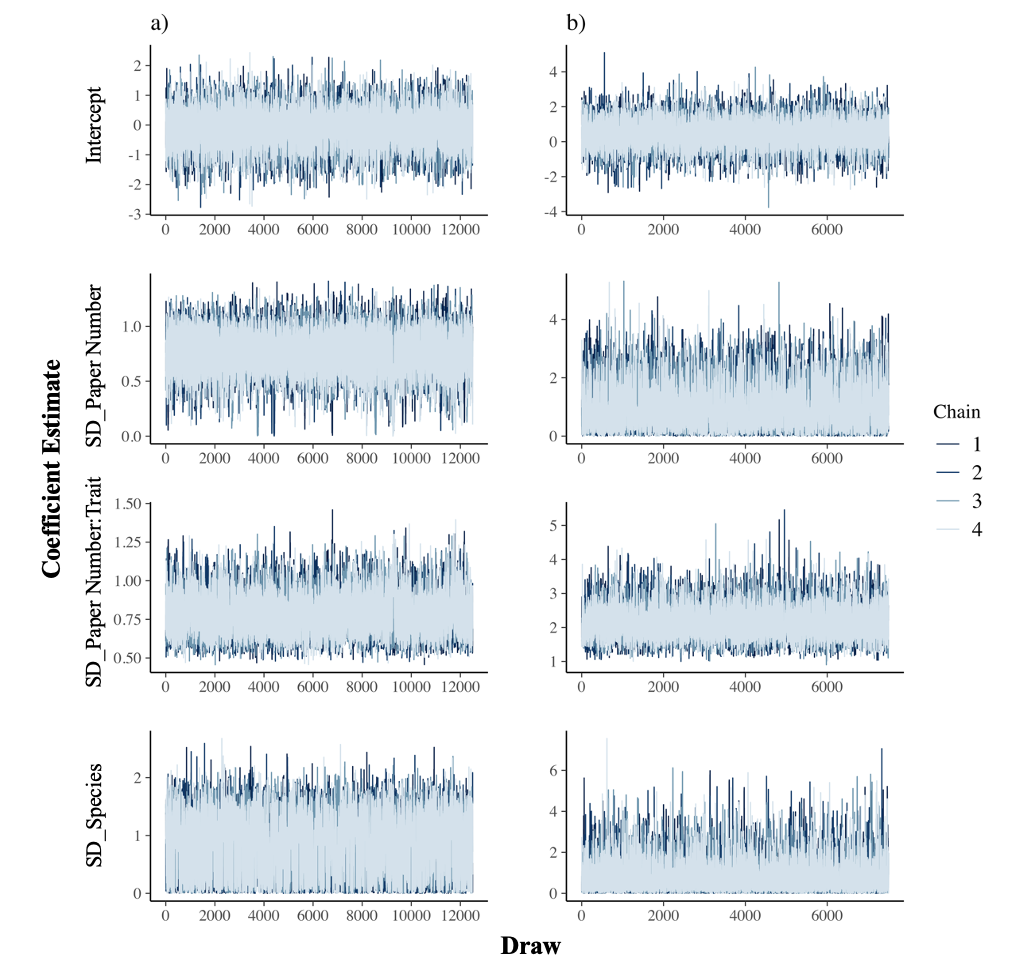

**Figure S2.** Trace plots for a) countergradient and b) cogradient random effects models. These plots illustrate that proper chain mixing occurred at each draw (x-axis) for the MCMC algorithm for each coefficient in the random effects models.

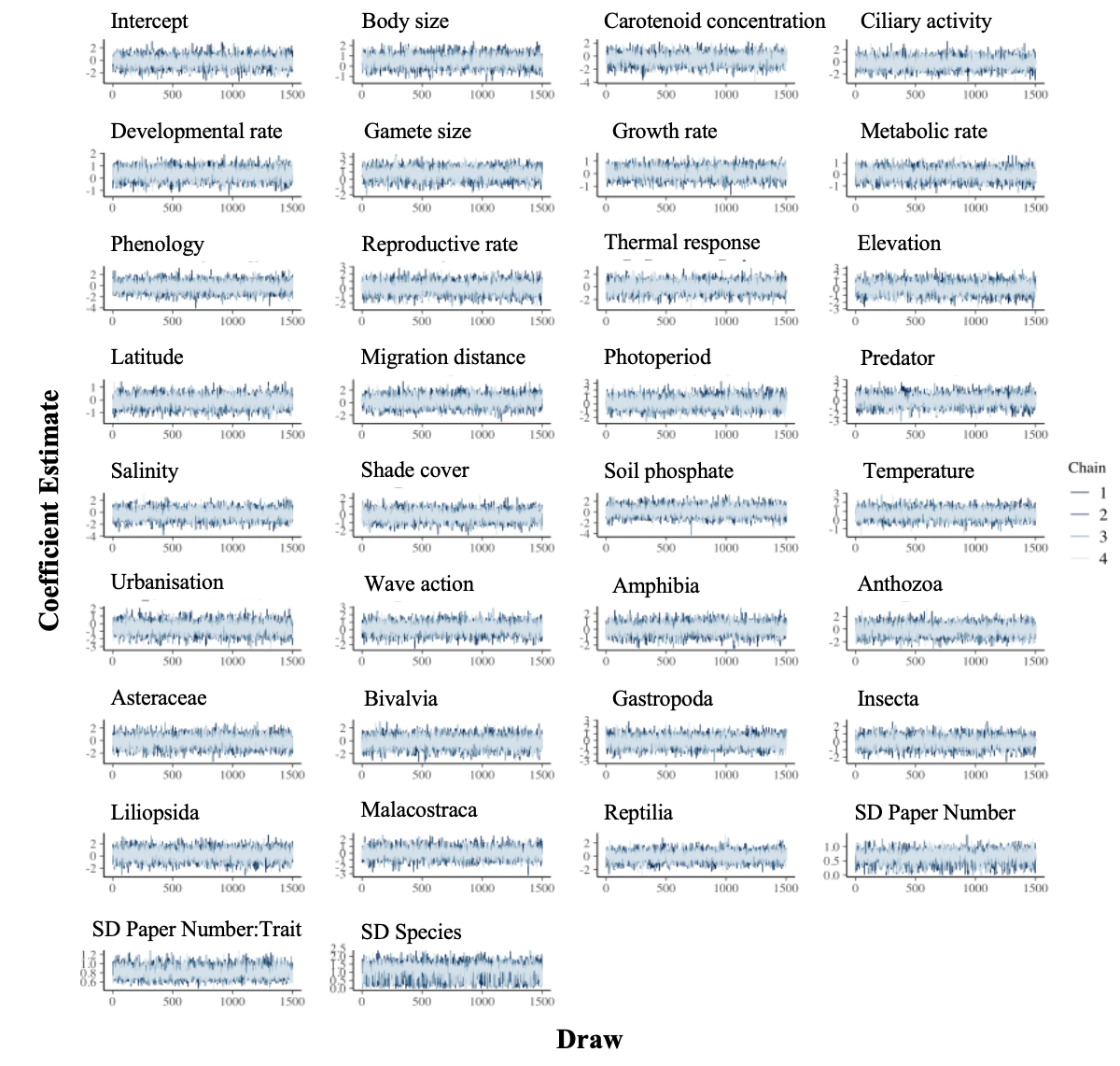

**Figure S3.** Trace plot for countergradient metaregression. These plots illustrate that proper chain mixing occurred at each draw (x-axis) for the MCMC algorithm for each coefficient in the metaregression model.

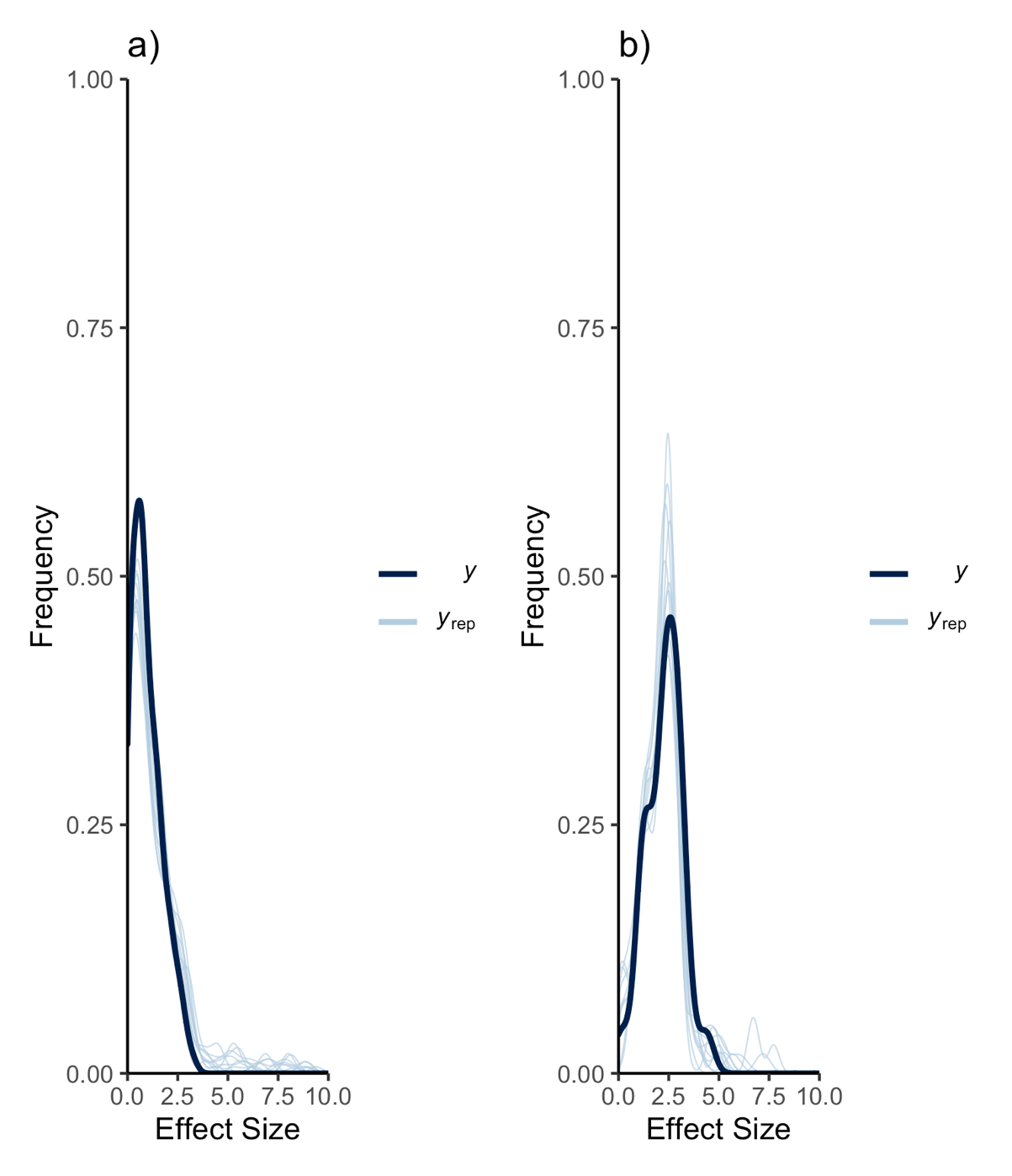

**Figure S4.** Posterior predictive checks for a) countergradient and b) cogradient random effects models. Posterior predictive checks are a tool to simulate replicated data and then to compare those simulations (light blue) with observed data (dark blue). The x-axis is the transformed effect size and the y-axis is the frequency.

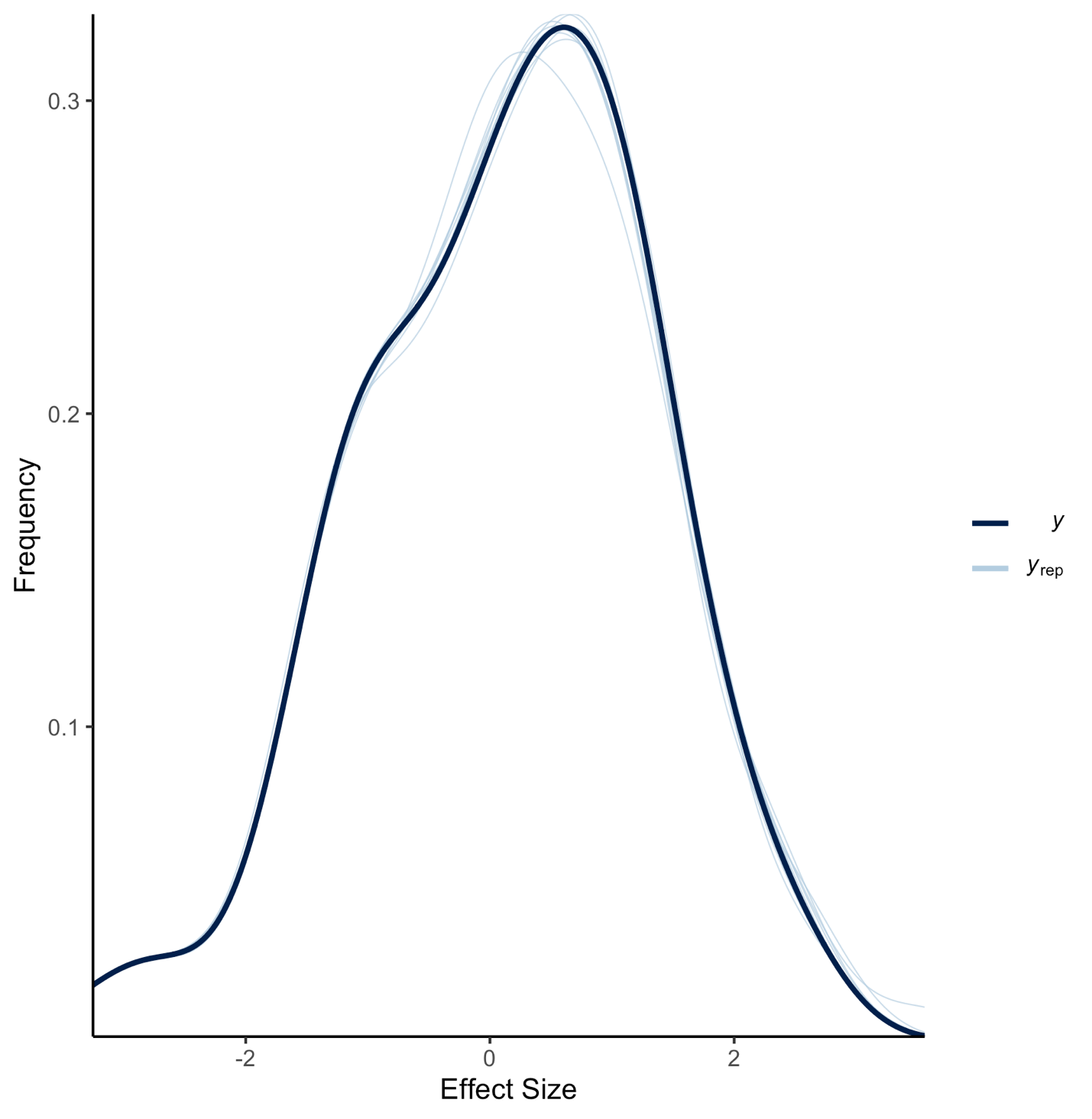

**Figure S5.** Posterior predictive checks for countergradient metaregression model. Posterior predictive checks are a tool to simulate replicated data and then to compare those simulations (light blue) with observed data (dark blue). The x-axis is untransformed effect size and the y-axis is the frequency.

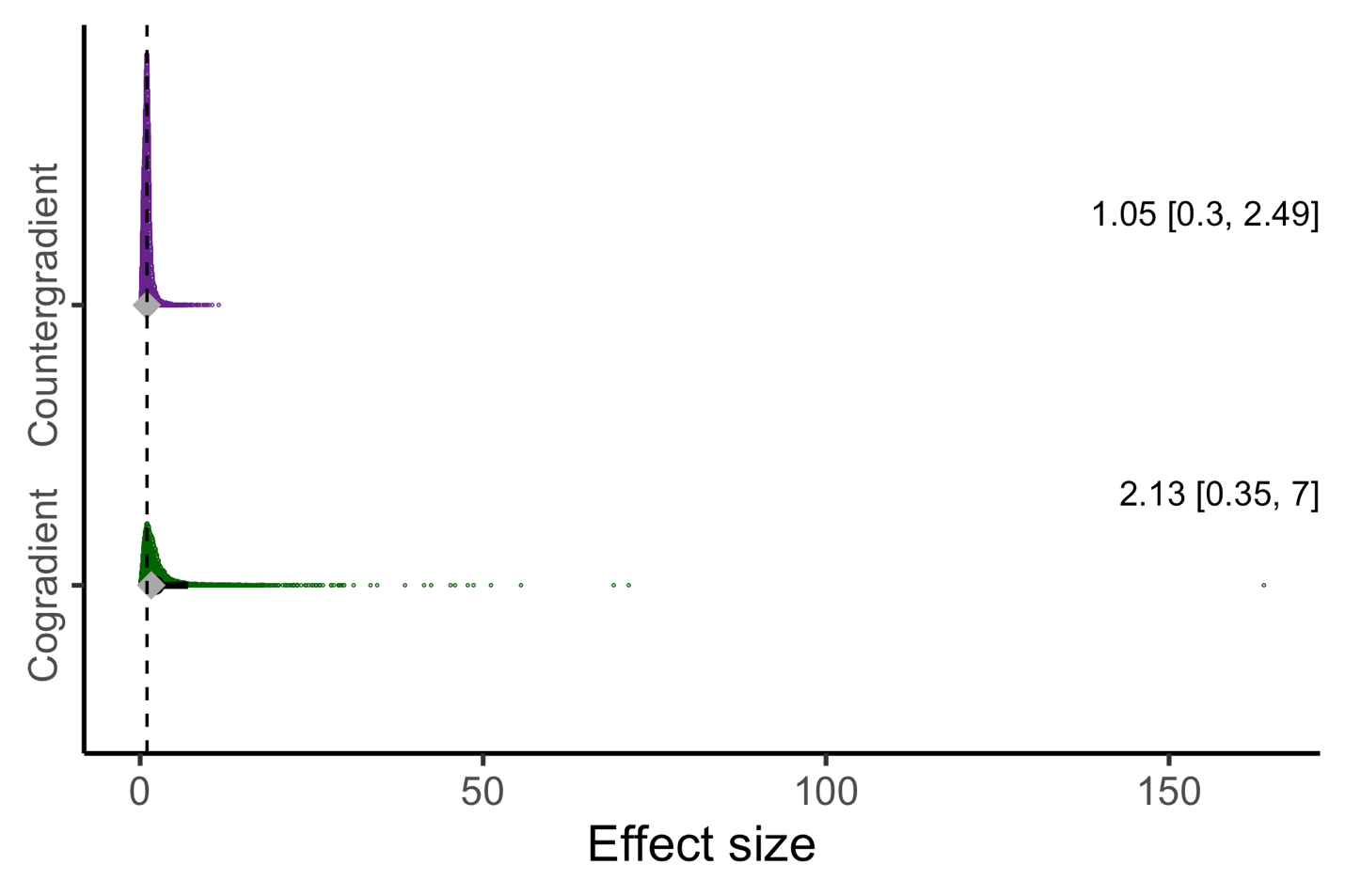
**Figure S6.** Posterior distributions of effect size estimates from random-effects meta-analysis for counter- and cogradient variation. The circle indicates the mean and the diamond indicates the median of the posterior distribution (overlapping) and the solid, black line, the 95% credible intervals. The mean and credible interval values are also indicated in text on the right. The dashed vertical line is meant to show the posterior distribution’s relationship to an effect size of 1. In this figure all data are shown (*c.f.* Fig. 2. where axis includes points between 0 and 8).

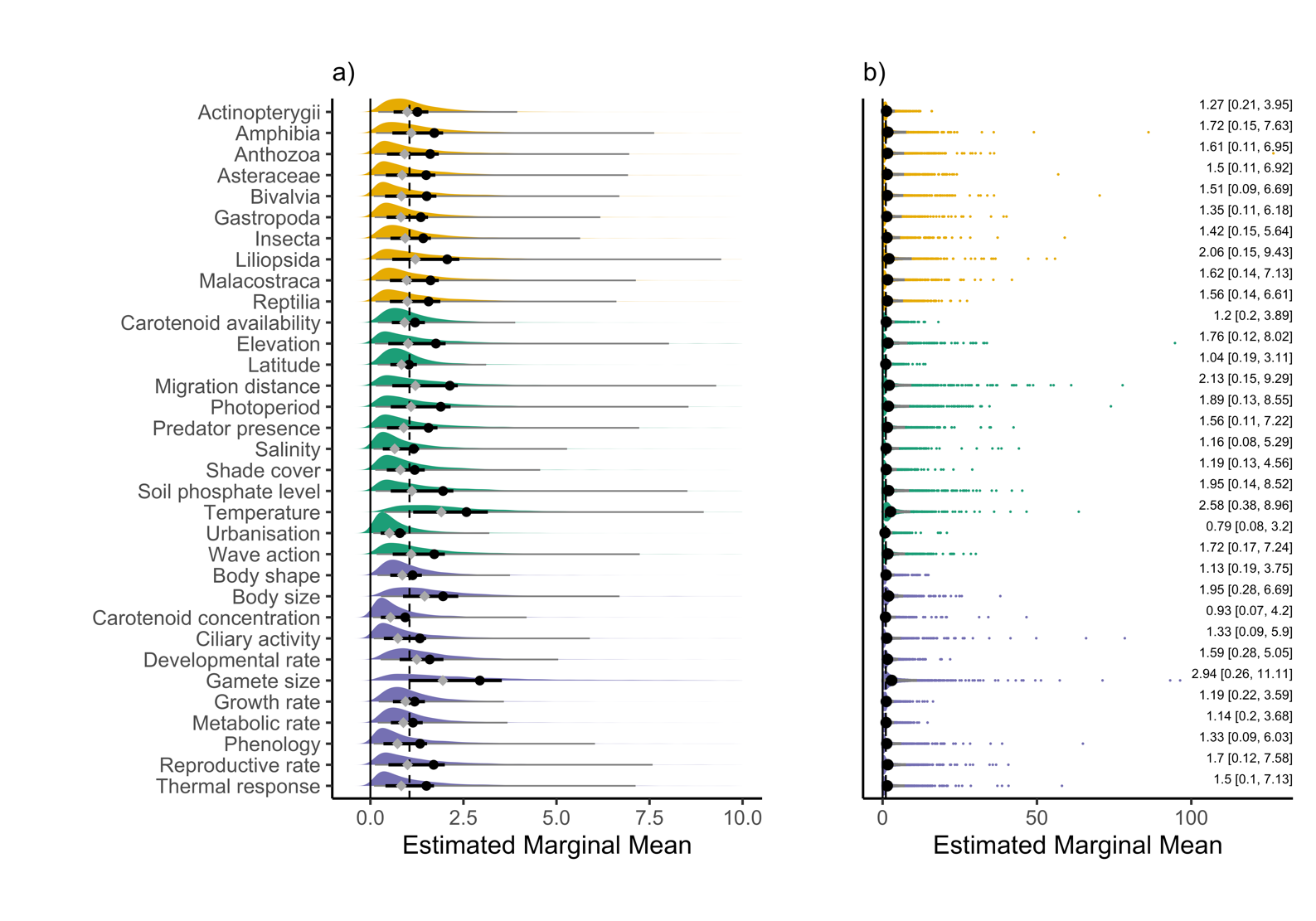

**Figure S7.** Main effects estimated marginal means for posterior distributions of covariates in metaregression analysis for countergradient variation. Yellow distributions are for organism’s class, green distributions are for the gradient covariate, purple for trait covariate. The circle indicates the mean the posterior distribution and the solid, black line, the 50% credible intervals and the thinner, grey line, the 95% credible intervals. The vertical dashed line illustrates the posterior distribution’s relationship to an effect size of 1.05 estimated from the random intercepts model and shown in Fig. 2 (*c.f.* Fig. 3. where axis includes points between -2 and 5). Panel a) is meant to illustrate the full 95% credible interval distribution, some of which are cut off in Fig. 3, and panel b) shows every value of the posterior distribution.

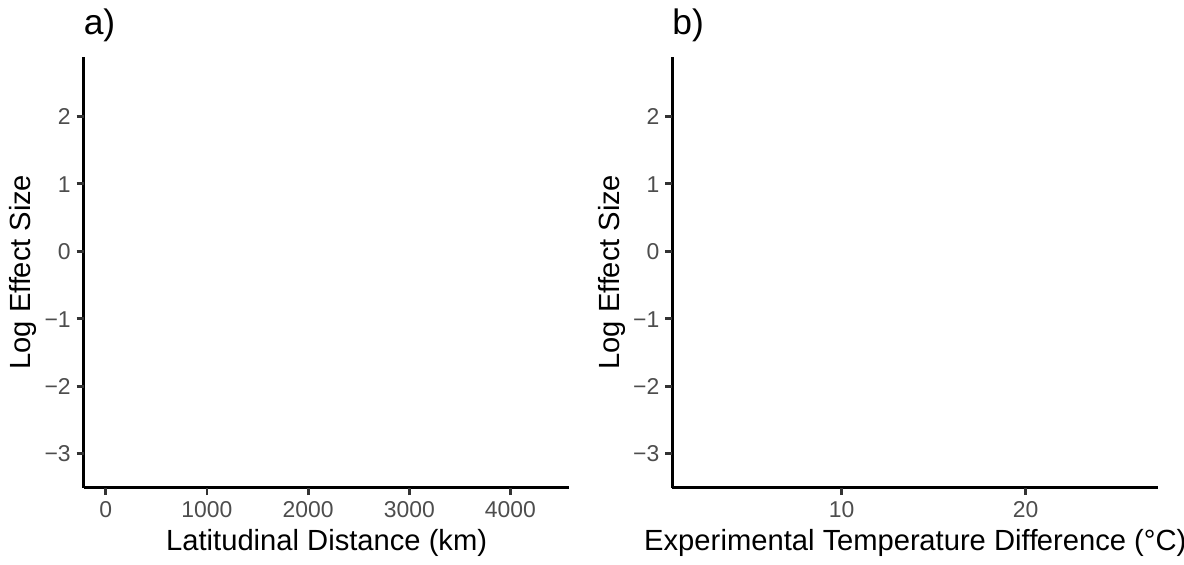

**Figure S8:** Effect size of countergradient variation as a function of a) latitudinal distance (km) between the most extremely distributed populations in a given study or difference in b) temperature (Celsius) in a given study. Neither represent a statistically significant effect (Table S7), nor were they predictive of effect size. Effect sizes are natural log transformed to address the non-normality and heteroscedasticity in the data.
